# Imbalanced classification for protein subcellular localisation with multilabel oversampling

**DOI:** 10.1101/2022.09.12.507675

**Authors:** Priyanka Rana, Arcot Sowmya, Erik Meijering, Yang Song

**Affiliations:** School of Computer Science and Engineering, University of New South Wales, Sydney, NSW 2052, Australia

## Abstract

**Motivation:** Subcellular localisation of human proteins is essential to comprehend their functions and roles in physiological processes, which in turn helps in diagnostic and prognostic studies of pathological conditions and impacts clinical decision making. Since proteins reside at multiple locations at the same time and few subcellular locations host far more proteins than other locations, the computational task for their subcellular localisation is to train a multilabel classifier while handling data imbalance. In imbalanced data, minority classes are underrepresented, thus leading to a heavy bias towards the majority classes and the degradation of predictive capability for the minority classes. Furthermore, data imbalance in multilabel settings is an even more complex problem due to the coexistence of majority and minority classes.

**Results:** Our studies reveal that based on the extent of concurrence of majority and minority classes, oversampling of minority samples through appropriate data augmentation techniques holds promising scope for boosting the classification performance for the minority classes. We measured the magnitude of data imbalance per class and the concurrence of majority and minority classes in the dataset. Based on the obtained values, we identified minority and medium classes, and a new oversampling method is proposed that includes nonlinear mixup, geometric and colour transformations for data augmentation and a sampling approach to prepare minibatches. Performance evaluation on the Human Protein Atlas Kaggle challenge dataset shows that the proposed method is capable of achieving better predictions for minority classes than existing methods.

**Availability:** Data used in this study is available at https://www.kaggle.com/competitions/human-protein-atlas-image-classification/data.

**Supplementary information:** Supplementary data are available at *Bioinformatics* online.

## 1 Introduction

A cell contains approximately 10^9^ proteins residing at different subcellular locations (Chou and Shen, 2006), namely organelles which are well defined membrane bound compartments for performing specific cellular functions. The subcellular location of a protein defines its functionality and helps understand protein behaviours. Any abnormal translocation of proteins in subcellular environments is vital to understand the clinical diagnosis and drug targeting treatment.

The spatial patterns of protein distribution in high contrast immunofluorescence images are interpretable, intuitive and can be effectively utilised for the detection of protein translocation. However, protein subcellular localisation (PSL) is challenging due to the proteins being located in multiple organelles. Additionally, proteins exhibit highly variant distribution, which makes PSL an imbalanced multilabel classification task. Accordingly, the classification model is expected to recognise protein patterns among different organelles and also cell phenotypes, which further adds to the complexity of PSL. More specifically, the model classifies protein images into multiple subcellular locations provided in the label sets. The difference in label frequencies across the dataset leads to varied representation of classes (subcellular locations/organelles) during training. The majority classes with higher number of samples are frequently observed during training, and minority classes with lower number of samples are rarely encountered. Subsequently, the trained model can be heavily biased towards the majority classes and demonstrate very poor performance on minority classes.

In the past, PSL has been studied extensively using handcrafted features (Xu *et al*., 2018; Rana *et al*., 2021). Advancements in microscopic imaging has led the focus of current studies towards deep learning with state-of-the-art performance. Existing deep learning based PSL methods have utilised different datasets and approaches to handle data imbalance. The Human Protein Atlas (HPA) project is the widely used public imaging database to study human proteins (Thul and Lindskog, 2018). In reference to the HPA project, the “Human Protein Atlas Image Classification” challenge is hosted on the Kaggle platform (Ouyang *et al*., 2019). The winning team utilised multilabel full-sized (a mix of 2, 048 × 2, 048 and 3, 072 × 3, 072 pixels) images from the Kaggle competition and external sources, and replaced the labels provided with protein IDs. Thereafter they applied metric learning using ArcFace loss (Deng *et al*., 2019). Another study (Zhang *et al*., 2021) utilised the same image sets, and applied threshold optimisation, cycle learning (Smith, 2017) and a weighted loss function with a hard-data sampler for improved performance. In order to save computational cost, Wang (2020) utilised only scaled Kaggle images (512 × 512 pixels) and proposed a novel composite loss function, which combines focal loss (Lin *et al*., 2017) and Lovász-Softmax loss (Berman *et al*., 2018). They trained three different models which were later ensembled to achieve better results. Aggarwal *et al*. (2022) proposed a convolutional neural networks (CNN) based model which outperformed the stacked ensemble of three pretrained CNN models.

Recently, PSL methods have incorporated multi-instance learning settings where labels are provided for each protein instead of an image, and each protein is considered a bag of multiple images/instances. Arcamone *et al*. (2021) applied few shot learning under which the model is trained on majority classes using a contrastive representation learning framework (Le-Khac *et al*., 2020) with ArcFace loss function and cosine distance metric. During testing, the trained feature extractor provides the feature embeddings for minority classes and evaluation is performed using a 9-way 20-shot learning scheme. Tu *et al*. (2022) proposed a new method, namely SIFLoc, which performs training in two stages. Stage one uses a contrastive learning based self-supervised pretraining method which includes hybrid data augmentation and a modified contrastive loss function, followed by stage two of supervised learning. Another PSL study by Zhang *et al*. (2022) applied multi-task learning strategy for pretraining and proposed a combination of discrimination loss and reconstruction loss for better performance. During prediction, a multi-instance method is applied with balanced sampling strategy.

These existing methods have mostly utilised different subsets of HPA that are different in the number of classes, images and degree of imbalance. They have achieved improvement over their base models; however, their performance with respect to each other is not known. Furthermore, all the methods so far have focussed on image feature description, to make the model learn to distinguish different classes. However, there are no existing methods that have been specifically designed for the problem of multilabel imbalanced classification in PSL.

Data imbalance is identified when certain classes have far fewer samples in the dataset than other classes. However, in multilabel settings, data imbalance is also in reference to label sets. A label set constitutes all the classes associated with a sample, which can contain different combinations of classes with various number of classes per sample, causing further data imbalance. Therefore, in order to handle the data imbalance in multilabel settings effectively, it is essential to quantify the imbalance, particularly the magnitude of concurrence among majority and minority classes. The SCUMBLE (Score of ConcUrrence among iMBalanced LabEls) metric (Charte *et al*., 2014) was proposed to assess the magnitude of concurrence in a multilabel dataset. Values for SCUMBLE fall in the range [0,1], where a value ≤ 0.1 indicates not much coexistence of minority and majority classes in the dataset and vice versa.

Existing techniques to handle multilabel data imbalance can be categorised into classifier dependent and independent approaches. Classifier dependent approaches include ensembling, algorithmic-specific adaptations and cost-sensitive learning (Tarekegn *et al*., 2021). Ensemble methods utilise different classifier models trained on the same data to obtain diverse predictions which are combined to boost the classifier performance. Algorithmic-specific adaptations enhance existing machine learning algorithms, for instance multilabel-KNN (Zhang and Zhou, 2007), multilabel SVM (Elisseeff and Weston, 2001) and multilabel neural networks (Zhang and Zhou, 2006; Zhang, 2009). Cost sensitive learning applies high cost on the misclassified minority samples. It is however not a widely applied method as it is challenging to determine an effective cost matrix for the dataset (Tarekegn *et al*., 2021).

Classifier independent approaches to handle class imbalance include oversampling of minority samples. However due to the coexistence of majority and minority classes in multilabel images, only the datasets with low SCUMBLE metric value benefit from oversampling techniques (Charte *et al*., 2015a, 2019a,b). Oversampling of multilabel samples can be performed by approaches such as LP-ROS (Label Powerset-Random Oversampling) (Charte *et al*., 2013) and ML-ROS (Multi Label-Random Oversampling) (Charte *et al*., 2015a). Both clone the original minority instances using various geometric and colour transformations (GCTs). LP-ROS considers the label set to evaluate imbalance while ML-ROS evaluates the imbalance level of each label/class. These methods if applied repetitively, often cause overfitting.

In order to create more variations in synthetic images, Multilabel Synthetic Minority Oversampling Technique (MLSMOTE) (Charte *et al*., 2015b) was proposed to create new images by interpolating two neighbouring images from the pool of minority class samples. However, generation of new instances and their corresponding labels in multilabel settings is often associated with chances of introducing noise alongside. Accordingly, Charte *et al*. (2019b) proposed a resampling approach for datasets with high concurrence, namely REMEDIAL (REsampling MultilabEl datasets by Decoupling highly ImbAlanced Labels), which decouples imbalanced concurrent labels and applies resampling for improved classification performance. The “mixup” method (Zhang *et al*., 2018) and its variants have been widely used to handle data imbalance in multiclass and single label settings (Chou *et al*., 2020; Galdran *et al*., 2021; Tarekegn *et al*., 2021; Shorten and Khoshgoftaar, 2019), however its application in multilabel settings has not been explored much.

In this study, we propose an oversampling method to handle data imbalance in multilabel settings, by creating synthetic samples using multiple data augmentation techniques combined with an imbalance aware sampling approach. In multilabel settings, each sample is associated with more than one class, due to which oversampling often generates noise. Therefore, it is crucial to identify the classes for oversampling and design the data augmentation approach accordingly. We first investigate the imbalance ratio per class and the extent of concurrence of majority and minority classes using the SCUMBLE metric. Based on the value obtained, the benefit of applying oversampling to handle data imbalance is measured. Thereafter, our oversampling method is applied to handle data imbalance in multilabel protein images for PSL.

The proposed oversampling method constructs synthetic samples using *nonlinear mixup* and random combinations of GCTs to achieve better diversity in the training set for improved classification performance. Our *nonlinear mixup* method enhances the regularisation capability of mixup by combining the transformations in the spatial domain and colour space. In particular, regularisation is further enhanced in the spatial domain by using a matrix as mixup coefficient, followed by applying 3D rotation in colour space about different reference channels which correspond to proteins of interest and the relevant cellular regions of minority classes. We also design an imbalance aware sampling approach, which measures imbalance using the median and mean of the “imbalance ratio per class” values and accordingly identifies the medium and minority classes. Thereafter, synthetic samples of minority and medium classes form separate minibatches for training and supplement the minibatch of original images during different iterations, so that the minority class samples are utilised more than the medium class samples.

The proposed oversampling method improves our previous approach (Rana *et al*., 2022a) that handles data imbalance in single label settings. The proposed improved method uses a matrix as a mixup coefficient instead of the standard scalar. Furthermore, *nonlinear mixup* images are generated by applying 3D rotation specifically about the colour components (image channels) that correspond to only minority classes, unlike our previous approach which randomly selects the axis of 3D rotation for single channel grey scale images. In addition, our new imbalance aware sampling approach measures the imbalance in the dataset and focusses on medium and minority classes, while our previously proposed minority focussed sampling approach did not include any such measures and focussed only on the minority classes. The proposed method was evaluated on the Kaggle challenge dataset (Ouyang *et al*., 2019) as it contains the highest number of classes among all the HPA subsets, and with the most extreme data imbalance closest to real world settings (Figure 1). The results demonstrate that our proposed framework achieves better predictions for minority and medium classes than existing methods.

**Fig. 1.**
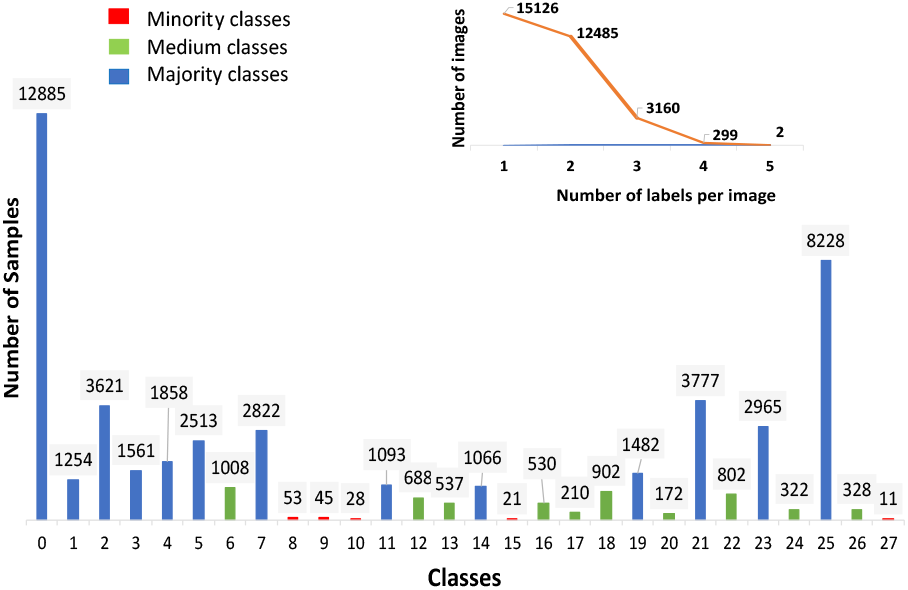
Class distribution of the Kaggle challenge dataset used in this study, with the number over the bar indicating the frequency of that class. Class numbers (0-5) represent the nuclear region and (6-27) represent the outer nuclear region, where (6-9) and (19-22) represent microtubules, while (10-18) and (23-27) represent endoplasmic reticulum. Each image has 1 to 5 labels.

## 2 Materials and methods

### 2.1 Dataset

The Kaggle challenge dataset has 28 classes, 31, 072 samples for training and 11, 702 samples for testing (TS2) (labels are not provided for the test set). Each sample consists of 4 channels: green, red, blue and yellow representing the antibody-stained protein of interest (POI), microtubules, nucleus and endoplasmic reticulum respectively (see Supplementary Figure 1). The goal of the study is to predict the subcellular location of POI, which is in the green channel, while red, yellow and blue channels function as references. Images are available in their original high-resolution size of 2, 048 × 2, 048 pixels and the downsized version of 512 × 512 pixels. The class distribution of the dataset with the number of samples in each class and number of images with their respective label set sizes are shown in Figure 1. There are 1 to 5 labels per image.

Due to the unavailability of test set labels, we randomly selected 20% of images from each class of the given training dataset to use as another test set (TS1) for the evaluation of methods.

### 2.2 Methods

First we measure the extent of concurrence of the majority and minority classes in the dataset to assess if oversampling can boost the classification performance. Next, minority and medium classes are identified (Figure 1) and our proposed oversampling approach is employed for model training.

#### 2.2.1 Data imbalance measures

Data imbalance in a multilabel dataset can be measured using two main measures, namely Imbalance Ratio per Label (IRL) and SCUMBLE (Tarekegn *et al*., 2021). In a multilabel dataset *M* with *L* labels/classes, *Y* label sets, *Y*_*i*_ denoting the label set of an instance *i*, IRL of a label *l* is the ratio of the number of samples of the most frequent label (majority class) to the label *l*:

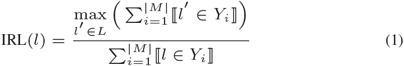

In addition, SCUMBLE measures the coexistence of majority and minority labels in the multilabel dataset as:

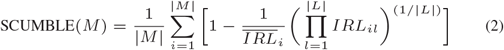

where IRL_*i*_=IRL(*l*) if label *l* is present in the instance *Y*_*i*_, else it is 0 and 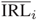 denotes the average imbalance level of the labels appearing in the *i*th instance. A low SCUMBLE value implies lower concurrence of imbalanced labels. A value ≤ 0.1 is taken to benefit from resampling (Charte *et al*., 2019b).

#### 2.2.2 mixup

In this study we used mixup as a data augmentation technique to improve the regularisation effect on the classifier in multilabel settings. It adopts the standard mixup strategy that generates synthetic images through weighted linear interpolation of two input images (Zhang *et al*., 2018):

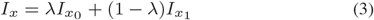

where 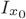 and 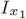 are two image matrices and *λ* is the mixup coefficient for each sample pair, which is a scalar ∈ [0, 1] in standard settings.

In order to obtain better regularisation, this study used *λ* as a matrix of random values ∈ [0, 1] which is of the same size as the image. Matrix *λ* provides more diverse variations in the resultant mixup image as it expands the space between reference images to generate synthetic images. In order to avoid the *mixup* image being too close to the reference images, the elements of mixing matrix *λ* are set to random values in the range of 0.35 to 0.65. A pair of images from the same minority class is randomly chosen and interpolation is carried out between the corresponding pixels of two reference images to generate a new mixup image. The label set of the new mixup image is the intersection of the label sets of the two reference images, where intersection refers to the labels which appear in both reference images.

#### 2.2.3 Nonlinear mixup

In order to enhance the regularisation effect of mixup, we earlier proposed a nonlinear mixup method (Rana *et al*., 2022a) to modify the pixel values of the mixup image in colour space using 3D rotation. Nonlinear mixup produces a synthetic image with transformations in both the spatial domain and colour space to provide better regularisation effects, compared to the standard mixup. Nonlinear mixup is carried out as follows: Step 1: For two randomly chosen reference images from a certain minority class, mixup is first used to create a new synthetic image *I*_*mix*_ through weighted interpolation of the corresponding channel images using matrix as mixup coefficient. This step transforms pixel values in the spatial domain. Step 2: Each pixel of *I*_*mix*_ obtained from Step 1 undergoes 3D rotation in colour space for further transformation. 3D rotation is carried out about one of the colour axes at a randomly chosen angle *θ* using the corresponding rotation matrix. Consequently, the pixel value of the colour component which is the axis of rotation remains unchanged, while new pixel values for the other two colour components are created to generate a different colour appearance. Step 3: The label set of the obtained nonlinear mixup image is inherited from its corresponding mixup image, which is the intersection of the label sets of the two input images.

To adapt this nonlinear mixup method to multilabel HPA images, consider that HPA images have four channels: green channel as POI, blue channel as the reference for the inner nuclear region, red and yellow channels as the references for the outer nuclear region. Since mixup and nonlinear mixup are applied only to the minority classes which are subcellular locations in the outer nuclear region, in this study the blue colour component of the 3D RGB colour space is replaced with the yellow colour component to construct a 3D colour space including red, green and yellow colour components (Figure 2). Subsequently, nonlinear mixup is used to generate two image variants, namely NL mixup (G) and NL mixup (R) by applying 3D rotation to the obtained mixup image about the green and red colour component respectively. 3D rotation about the green axis keeps the POI pixels unchanged, while creating variations in the outer nuclear region. Consequently, NL mixup (G) has the same POI pattern as mixup but with variant backgrounds (Figure 2). NL mixup (R) brings variations in the green and yellow colour components while the red colour component remains unchanged. With a controlled tweak such as setting up the range to select *θ*, NL mixup (R) accentuates POI pixels (Figure 2). More specifically, each pixel in the 3D colour space represented as 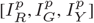 is rotated at an angle *θ* using rotation matrices *Rot*_*G*_ and *Rot*_*R*_ for NL mixup (*G*) and NL mixup (R) respectively (Figure 2). During 3D rotation about *G*, a new pixel value is created on a circle whose centre is 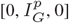 with radius 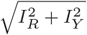, and similarly for the rotation about *R*. Accordingly, transformation of pixels for NL mixup (G) and NL mixup (R) are defined as:

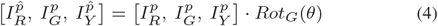

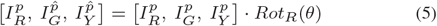

respectively, where *Rot*_*G*_ and *Rot*_*R*_ are:

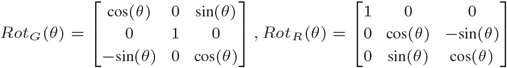

In order to obtain more variations in the nonlinear mixup images across the iterations and avoid the resultant NL mixup image being too similar to the mixup image, *θ* is randomly chosen alternatively from two sets ([60^°^- 110^°^], [120^°^- 300^°^]).

**Fig. 2.**
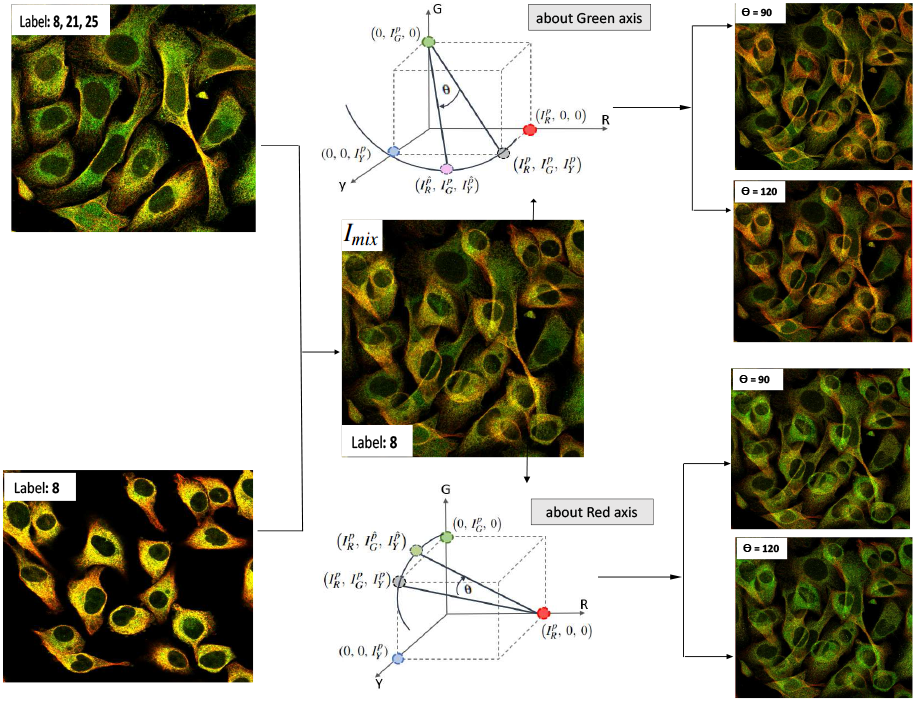
Mixup and nonlinear mixup (RGY view): Two images randomly selected from the same minority class ‘8’ to generate a mixup image (*I*_*mix*_) with a label obtained from the intersection of the label sets of the reference images. Nonlinear mixup image is generated by applying 3D rotation to each pixel about the green and red axes in the colour space. Rotation about the green axis at two different angles *θ* provides variation in backgrounds to POI (green channel). Rotation about the red axis at two different angles *θ* accentuates POI pixels.

#### 2.2.4 Imbalance aware sampling

The standard approach to identify classes that need oversampling is to consider all the classes in the dataset with IRL *>* mean 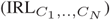 as minority classes. However, in cases of extreme data imbalance, the distribution of IRL values of the classes is skewed to the right. Therefore, apart from minority classes, there is another category of classes that may benefit from oversampling, that we have named medium classes. The medium classes have IRL too high to be considered as majority class and too low to be a minority class. Since for a right skewed distribution, the mean is typically greater than the median, this study utilises the median of the IRL values to identify medium classes. Accordingly, all the classes which satisfy *median* < IRL < *mean* are considered to be medium classes.

Adapting from our earlier approach (Rana *et al*., 2022b), in this study, each minibatch during an iteration is supplemented with another minibatch of synthetic samples from minority or medium classes. Our sampling approach does not require a preset sample distribution ratio, instead it finetunes the number of samples per class to be added for training during an iteration through an hyperparameter *n*. For the supplementary batch of minority samples, *n*=1 adds three images (original image (with GCT) + NL mixup (R) + NL mixup (G)), while the supplementary batch of medium samples consists of all three images obtained after random combinations of GCTs. Consequently, the method balances the representation of target classes at minibatch level and reduces computational cost as it allows the inclusion of the minimum number of synthetic samples, without affecting the representation of the original samples.

We note that 65% of the iterations per epoch includes the supplementary batch of minority samples and 35% of iterations per epoch includes the supplementary batch of medium samples. Since the number of original samples in medium classes are comparatively high with a wide range of label set sizes, it is observed that if mixup/NL mixup is applied to medium classes, the new label sets obtained after intersection affects the predictions of other classes. Therefore, only GCTs are applied on medium class samples during a small percentage of iterations.

## 3 Experimental setup and implementation

The proposed method and several state-of-the-art PSL methods were evaluated on the multilabel HPA Kaggle challenge images of size 512×512 pixels. We computed IRL and SCUMBLE metrics to measure the data imbalance and, based on the IRL values, identified the minority and medium classes. Minority classes are 8, 9, 10, 15, 27, and medium classes and therefore oversampling could be useful to handle the data imbalance. Accordingly, minority samples were constructed using GCTs and mixup, which were further transformed using nonlinear mixup, while oversampling of medium classes was performed using only GCTs. We utilised GCTs such as horizontal/vertical/left/right flips, rotation at 90^°^, 180^°^, 270^°^, brightness adjustments of images in the range of 50-150% of the original pixel value and cropping followed by padding.

For classification, the available training data after separation of TS1 images was divided as follows: 80% for training, and 20% for validation, for 5-fold stratified cross-validation. Subsequently, five models were obtained after training on each set of training and validation images. The final predictions for TS1 and TS2 images was obtained by ensembling via averaging the predictions of all five models. The classification model was trained by finetuning the ImageNet pretrained ResNet-50 (He *et al*., 2016) architecture on the training dataset, which is a common model used for PSL studies. The loss function was binary cross entropy (BCE). The minibatch size was 32, where the number of original samples was 17 and *n* was set to 1 for all the minority classes. The weight parameters were updated using the stochastic gradient descent method (Robbins and Monro, 1951) with a momentum parameter of 0.9 and weight decay of 0.0001 over 60 epochs. The initial learning rate was 0.03 and StepLR offered by PyTorch was used to schedule the learning rate, with gamma and step size set to 0.05 and 15 respectively. The initial settings of hyperparameters (batch size, decay and learning rate) and learning scheduler, were the same for all the experiments.

Hyperparameter temperature for SIFLoc (Tu *et al*., 2022) was 0.1. This study used the macro F1-score for performance evaluation of the proposed method, which is the arithmetic mean of the F1-scores of all 28 classes and indicates the performance of all the classes in the dataset equitably. Methods were evaluated based on the F1-scores of minority classes in the utilised test set (TS1), macro F1-score of TS1 and Kaggle test set (TS2). In addition, Wilcoxon rank sum test at significance level of 1% is used to demonstrate the statistical significance of the results. The Wilcoxon rank sum test compares the probabilities of class samples in TS1 obtained from the proposed method and other compared methods.

## 4 Results and discussion

We first compare the performance of the proposed method, base model and existing PSL methods which are based on contrastive learning (Tu *et al*., 2022), metric learning (Ouyang *et al*., 2019) and supervised learning using Focal-Lovász (Wang, 2020) loss function. The base model results are from the pretrained ResNet-50 model finetuned on the original images of the Kaggle challenge dataset using the BCE loss function.

As the results show (Table 1), our proposed method outperforms other compared methods. In reference to medium classes, our method improved their macro-F1 score from 0.61 to 0.66. As the IRL value of a class rises from the median value of IRLs, which is from class 6 to class 20 (see Supplementary Table 1), the percentage of improvement in the F1-score after oversampling also rises from 4.6% to 31.3% (see Supplementary Table 2).

**Table 1.** Comparison of PSL methods evaluated on the HPA Kaggle dataset. Best performance per class and per method is indicated in bold

| Study | C8 | C9 | C10 | C15 | C27 | TS1 | TS2 | p-value |
| --- | --- | --- | --- | --- | --- | --- | --- | --- |
| Our method | **0.82** | **0.88** | **1** | **0.57** | **0.67** | **0.74** | **0.478** |  |
| Base model (ResNet-50) | 0.62 | 0.75 | 0.70 | 0 | 0 | 0.65 | 0.431 | 8.7e-07 |
| Focal + Lovász loss (Wang, 2020) | 0.71 | 0.87 | 0.91 | 0 | 0 | 0.68 | 0.447 | 6.6e-06 |
| SIFLoc (Tu *et al.*, 2022) | 0.57 | 0.67 | 0.62 | 0.46 | 0 | 0.44 | 0.186 | 1.1e-46 |
| Metric learning (Ding *et al.*, 2015) | 0.20 | 0.16 | 0 | 0 | 0 | 0.32 | 0.224 | 1.5e-50 |

C8, C9, C10, C15, C27 are the F1-scores of minority classes 8, 9, 10, 15, 27 respectively and TS1 is the macro F1-score of the sampled test set (extracted from the Kaggle training set) in this study. TS2 is the macro F1-score of the official Kaggle test set obtained by submitting classification results to the Kaggle competition. F1-score for all individual classes in TS1 are shown in Supplementary Table 3.

In multilabel settings, since each sample is associated with more than one class, oversampling often generates noise due to a mismatch of the assigned label set to the newly constructed synthetic sample. Therefore, it is important to correctly identify the classes for oversampling before applying data augmentation. Our imbalance aware sampling approach measures imbalance in the dataset using mean and median of the “imbalance ratio per class” values and accordingly identifies the medium and minority classes. Noise and overfitting is avoided while handling imbalance, as synthetic medium samples are generated using only GCTs and supplement the original minibatch for fewer iterations, while synthetic minority samples are generated using NL mixup and GCTs, and supplement the original minibatch for majority of iterations. Noise is further avoided during NL mixup by selecting image pairs from the same class and reference channels that correspond to only minority classes, along with the intersection approach to generate new label sets. Consequently, our proposed method improves the generalisation ability of the model and makes it perform better on minority classes. For instance, as shown in Supplementary Figure 3, sample images have both majority and minority classes in their original label set, and Wang (2020) predicted only majority classes while our method is able to predict the complete label set correctly.

Among the compared approaches, SIFLoc (Tu *et al*., 2022) utilises a modified contrastive loss function to perform self-supervised pretraining followed by supervised learning. The model aims to capture the semantic relationships between images to generate better image representations. Moreover, unlike existing PSL methods that applied metric learning with ArcFace loss and protein IDs (Ouyang *et al*., 2019; Le-Khac *et al*., 2020), for further comparison, we also applied triplet margin loss function (Ding *et al*., 2015) to evaluate metric learning with multi labels (see Supplementary section 2 for details). Metric learning based on the triplet margin loss function does not perform well in comparison to other methods specifically for minority and medium classes. It could not capture the intraclass variance of minority samples as the chance of multiple minority class samples to fall together in a minibatch is low.

As shown in Table 1, supervised learning methods using BCE and Focal-Lovász loss perform better than SIFLoc and metric learning. However, the latter methods performed well in their respective PSL studies because of comparatively less extreme data imbalance in the utilised datasets and the use of protein IDs. In line with the existing study (Wang, 2020), Focal-Lovász loss achieves more than 3.5% improvement over the base model (with BCE loss) on both test sets in this study, however it does not perform as well as the proposed method.

For ablation studies of each component of the proposed approach, we explored different combinations of the utilised data augmentation techniques (GCTs, nonlinear mixup about red and green axis) and alternatives for nonlinear mixup such as standard mixup, MLSMOTE (Charte *et al*., 2015b) and manifold mixup (Verma *et al*., 2019). The compared methods are as follows: 1) GCT + mixup + NL mixup (G), which is oversampling of minority classes with images obtained after applying a random combination of GCTs, mixup and nonlinear mixup about the green axis, using imbalance aware sampling; 2) GCT + mixup + NL mixup (R), which is oversampling of minority classes with images obtained after applying a random combination of GCTs, mixup and nonlinear mixup about the red axis, using imbalance aware sampling; 3) mixup + NL mixup (R) + NL mixup (G), which is oversampling of minority classes with images obtained after applying mixup and nonlinear mixup about green and red axes, using imbalance aware sampling; 4) GCT + mixup, which is oversampling of minority classes with images obtained after applying a random combination of GCTs and mixup, using imbalance aware sampling; 5) GCT + manifold mixup, which is oversampling of minority classes with images obtained after applying a random combination of GCTs and manifold mixup, using imbalance aware sampling; 6) GCT + MLSMOTE, which is oversampling of minority classes with images obtained after applying a random combination of GCTs and MLSMOTE, using imbalance aware sampling; 7) GCT, which is oversampling of minority classes with images obtained after applying only a random combination of GCTs, using imbalance aware sampling; weighted sampling, which replaces imbalance aware sampling in the base model, assigns a weight that is the inverse of the class frequency to the samples during training and applies a random combination of GCTs; standard sampling, which replaces imbalance aware sampling and creates minibatches using a standard sampling strategy which randomly selects the images. The method applies data augmentation and performs oversampling of minority classes whenever it comes across a minority class sample. 10) scalar mixup coefficient, which replaces the proposed matrix mixup coefficient with a scalar mixup coefficient. We note that the model architecture, loss function, learning scheduler, optimiser and other initial hyperparameter settings were the same for all the experiments.

As shown in Table 2, the proposed oversampling framework that includes GCT, NL mixup (R) and NL mixup (G) with imbalance aware sampling achieves best performance in the ablation study. It is observed that both nonlinear image variants (NL mixup (R) + NL mixup (G)) contribute most to the performance of the classification model. Since NL mixup (R) accentuates POI pixels, while NL mixup (G) provides variations in the background of POI, this implies that the transformation of selective channels according to the task requirements with controlled tweaks can further enhance the regularisation effect. We note that, using a matrix as mixup coefficient improves the performance of the model by approximately 2% compared to using a scalar value.

**Table 2.** Ablation study.

| Study | C8 | C9 | C10 | C15 | C27 | TS1 | TS2 | p-value |
| --- | --- | --- | --- | --- | --- | --- | --- | --- |
| GCT + NL mixup (R) + NL mixup (G) | **0.82** | **0.88** | **1** | **0.57** | **0.67** | **0.74** | **0.478** |  |
| GCT + mixup + NL mixup (G) | 0.70 | 0.87 | 0.91 | 0.33 | 0.67 | 0.72 | 0.465 | 5.4e-03 |
| GCT + mixup + NL mixup (R) | 0.80 | 0.78 | 0.73 | 0.29 | 0.67 | 0.71 | 0.468 | 9.2e-04 |
| mixup + NL mixup (R) + NL mixup (G) | 0.78 | 0.87 | 0.91 | 0 | 0.67 | 0.71 | 0.450 | 7.1e-04 |
| GCT + mixup | 0.78 | 0.87 | 0.91 | 0.25 | 0.67 | 0.72 | 0.442 | 4.1e-03 |
| GCT + Manifold mixup | 0.67 | 0.80 | 0.91 | 0.25 | 0.67 | 0.69 | 0.440 | 1.2e-11 |
| GCT + MLSMOTE | 0.68 | 0.67 | 0.80 | 0 | 0 | 0.66 | 0.415 | 2.1e-19 |
| GCT | 0.70 | 0.80 | 0.77 | 0.20 | 0.50 | 0.70 | 0.442 | 3.3e-04 |
| Weighted Sampling | 0.73 | 0.87 | 0.83 | 0.40 | 0 | 0.68 | 0.452 | 1.4e-16 |
| Standard Sampling | 0.71 | 0.78 | 0.66 | 0 | 0 | 0.67 | 0.437 | 1.6e-18 |
| Scalar mixup coefficient | 0.78 | 0.83 | 0.84 | 0.38 | 0.67 | 0.72 | 0.467 | 6.3e-03 |

GCT stands for Geometric and Colour Transformations, NL mixup (G) and NL mixup (R) stands for nonlinear mixup with green and red colour component as axis for 3D rotation respectively. p-values smaller than 0.01 implies statistically improved performance of the proposed method over the compared methods.

The gap in the obtained macro F1-scores for TS1 and TS2 implies that TS2 contains images that are vastly different from the training images and therefore challenging to classify. It is observed that methods using two different data augmentation techniques do not show the same degree of performance improvement on TS2 (≈ 2% increase from the base model) as on TS1 (≈ 6% increase from the base model). On the contrary, more diverse data generated using three different data augmentation approaches, lead to considerable improvement for both TS1 (≈ 9.2% increase from the base model) and TS2 (≈ 9% increase from the base model). Experiments also demonstrated better performance with the imbalance aware sampling approach than standard sampling and weighted sampling approach. However, for the best results the proposed sampling approach must be applied with multiple data augmentation techniques to avoid overfitting.

Furthermore, as shown in Table 2, the proposed method with NL mixup (R/G) performs better than the standard mixup, MLSMOTE or manifold mixup. MLSMOTE applies interpolation to the image pair selected from the pool of the minority class samples using nearest neighbour approach without considering their respective classes. Since the dataset has images from different cell lines (see Supplementary Figure 2), a few synthetic images obtained from MLSMOTE contributed to the noise and exhibited comparatively poor performance. Manifold mixup applies interpolations in deep hidden layers between images from the same class for smoother decision boundaries. In current settings when applied on minority classes it affects the predictions of concurrent classes such as class 4, 6 and 19 reducing the overall macro F1 scores, however its performance on TS2 is almost as good as standard mixup.

As shown in Table 1 and Table 2, the obtained p-values are smaller than 0.01, which implies that the proposed method has statistically more significant performance than the compared methods.

## 5 Conclusion

In this paper, we have proposed an oversampling method to handle multilabel data imbalance for the protein subcellular localisation task. The method computes the value of the SCUMBLE metric which is used to determine the application of oversampling methods to handle data imbalance. Synthetic samples for oversampling are generated from *nonlinear mixup* and random combinations of geometric and colour transformations. The regularisation capability of *nonlinear mixup* is enhanced by using a matrix as the mixup coefficient, and applying 3D rotation in colour space about different reference channels which correspond to the protein of interest and the relevant cellular regions of minority classes. Synthetic samples are included for training through an imbalance aware sampling approach, which measures the imbalance ratio per class and accordingly identifies medium and minority classes. The proposed method evaluated on a publicly available Kaggle challenge dataset outperforms existing protein subcellular localisation methods. Experimental results demonstrate the advantage of applying oversampling using multiple data augmentation techniques and imbalance aware sampling approach to improve the predictions for minority and medium classes.

## Acknowledgements

This research was undertaken with the assistance of resources and services from the National Computational Infrastructure (NCI), which is supported by the Australian Government.

