## Supplementary materials for "Imbalanced classification for protein subcellular localisation with multilabel oversampling"

*Bioinformatics*
doi.10.1093/bioinformatics/xxxxxx
Advance Access Publication Date: Day Month Year
Manuscript Category

OXFORD

Bioimage informatics

### Imbalanced classification for protein subcellular localisation with multilabel oversampling

**Priyanka Rana, Arcot Sowmya, Erik Meijering, Yang Song** *

[1] School of Computer Science and Engineering, University of New South Wales, Sydney, NSW 2052, Australia

* To whom correspondence should be addressed.

Associate Editor: XXXXXXX

Received on XXXXX; revised on XXXXX; accepted on XXXXX

**Contact:**

#### 1 Supplementary tables and figures

Table 1. Class details with its corresponding imbalance ratio per label value

| Minority/Medium/Majority | Class Number (C) | Subcellular Localisation | Imbalance Ratio per label |
|---|---|---|---|
| **Minority** | 27 | Rods and rings | 1171.36 |
| **Minority** | 15 | Microtubule ends | 613.57 |
| **Minority** | 10 | Lysosomes | 460.18 |
| **Minority** | 9 | Endosomes | 286.33 |
| **Minority** | 8 | Peroxisomes | 243.11 |
| *Medium* | 20 | Lipid droplets | 74.91 |
| *Medium* | 17 | Mitotic spindle | 40.02 |
| *Medium* | 24 | Aggresome | 39.28 |
| *Medium* | 26 | Cytoplasmic bodies | 24.31 |
| *Medium* | 16 | Cytokinetic bridge | 24.0 |
| *Medium* | 13 | Focal adhesion sites | 18.73 |
| *Medium* | 12 | Actin filaments | 16.07 |
| *Medium* | 22 | Cell junctions | 14.29 |
| *Medium* | 18 | Microtubule organizing center | 12.80 |
| *Medium* | 6 | Endoplasmic reticulum | 12.0 |
| Majority | 14 | Microtubules | 11.77 |
| Majority | 11 | Intermediate filaments | 10.28 |
| Majority | 19 | Centrosome | 8.69 |
| Majority | 3 | Nucleoli fibrillar center | 8.25 |
| Majority | 4 | Nuclear speckles | 6.93 |
| Majority | 5 | Nuclear bodies | 5.13 |
| Majority | 7 | Golgi apparatus | 4.56 |
| Majority | 23 | Mitochondria | 4.35 |
| Majority | 2 | Nucleoli | 3.556 |
| Majority | 21 | Plasma membrane | 3.41 |
| Majority | 25 | Cytosol | 1.57 |
| Majority | 0 | Nucleoplasm | 1 |

.

© The Author 2015. Published by Oxford University Press. All rights reserved. For permissions, please 

Table 2. Improvement in F1-scores of medium classes after oversampling

| | **F1-score** | | |
|---|---|---|---|
| Class Number (C) | Base Model | Proposed method | %age increase |
| 20 | 0.48 | 0.63 | 31.3 |
| 17 | 0.58 | 0.64 | 10.3 |
| 24 | 0.74 | 0.80 | 8.1 |
| 26 | 0.48 | 0.54 | 12.5 |
| 16 | 0.50 | 0.54 | 8.0 |
| 13 | 0.67 | 0.70 | 4.5 |
| 12 | 0.71 | 0.74 | 4.2 |
| 22 | 0.65 | 0.68 | 4.6 |
| 18 | 0.63 | 0.65 | 3.2 |
| 6 | 0.65 | 0.68 | 4.6 |
| | 0.61 | 0.66 | |

.

Table 3. F1-score for all individual classes in TS1. Best performance per class and per method is indicated in bold.

| Class Number | Proposed method | Base model | Focal-Lovász | SIFLoc | Metric learning |
|---|---|---|---|---|---|
| 0 | **0.87** | 0.83 | 0.84 | 0.81 | 0.69 |
| 1 | **0.85** | 0.84 | 0.84 | 0.79 | 0.64 |
| 2 | **0.82** | 0.80 | 0.81 | 0.66 | 0.47 |
| 3 | **0.78** | 0.77 | 0.77 | 0.71 | 0.61 |
| 4 | **0.81** | 0.79 | 0.80 | 0.70 | 0.58 |
| 5 | **0.72** | 0.70 | 0.71 | 0.45 | 0.47 |
| 6 | **0.68** | 0.65 | 0.64 | 0.38 | 0.29 |
| 7 | **0.84** | 0.83 | 0.83 | 0.62 | 0.51 |
| 8 | **0.82** | 0.62 | 0.71 | 0.57 | 0.20 |
| 9 | **0.88** | 0.75 | 0.87 | 0.67 | 0.16 |
| 10 | **1.0** | 0.70 | 0.91 | 0.62 | 0.0 |
| 11 | **0.77** | 0.74 | 0.76 | 0.46 | 0.58 |
| 12 | **0.74** | 0.71 | 0.73 | 0.48 | 0.19 |
| 13 | **0.70** | 0.67 | 0.68 | 0.49 | 0.14 |
| 14 | **0.86** | 0.85 | **0.86** | 0.43 | 0.57 |
| 15 | **0.57** | 0.0 | 0.0 | 0.46 | 0.0 |
| 16 | **0.54** | 0.50 | 0.53 | 0.45 | 0.29 |
| 17 | **0.64** | 0.58 | 0.61 | 0.61 | 0.14 |
| 18 | **0.65** | 0.63 | 0.62 | 0.10 | 0.13 |
| 19 | **0.69** | 0.68 | 0.67 | 0.44 | 0.46 |
| 20 | **0.63** | 0.48 | 0.62 | 0.08 | 0.0 |
| 21 | **0.71** | 0.69 | 0.69 | 0.39 | 0.47 |
| 22 | **0.68** | 0.65 | 0.65 | 0.31 | 0.17 |
| 23 | **0.80** | **0.80** | 0.79 | 0.36 | 0.38 |
| 24 | **0.80** | 0.74 | 0.76 | 0.0 | 0.15 |
| 25 | **0.69** | 0.65 | 0.67 | 0.27 | 0.56 |
| 26 | **0.54** | 0.48 | 0.53 | 0.0 | 0.0 |
| 27 | **0.67** | 0.0 | 0.0 | 0.0 | 0.0 |
| **Macro F1-score** | **0.74** | 0.65 | 0.68 | 0.44 | 0.32 |

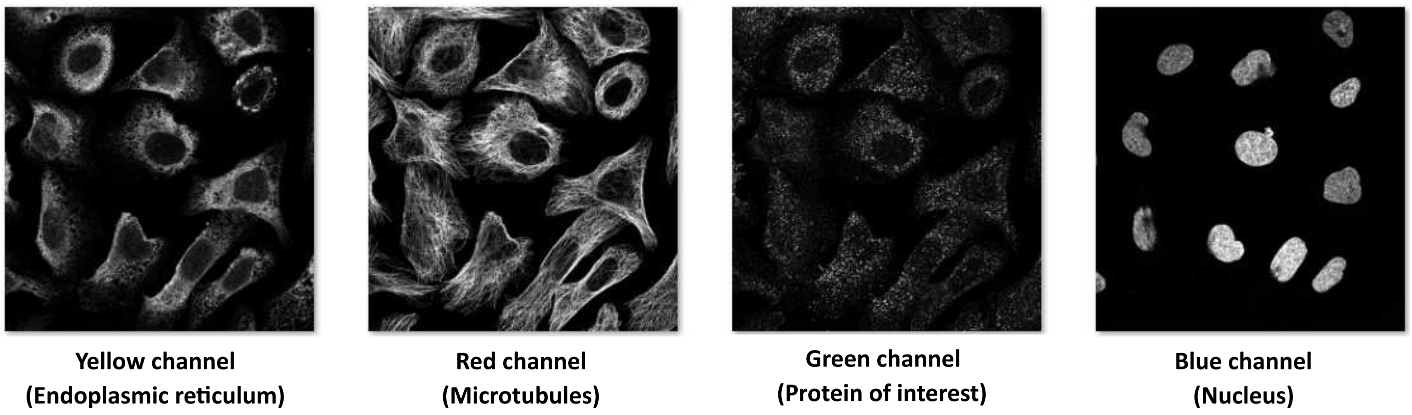


**Fig. 1.** Each sample consists of four channels: yellow, red, green and blue representing endoplasmic reticulum, microtubules, protein of interest (POI) and nucleus respectively. In this figure POI is class 8 which is Peroxisomes.

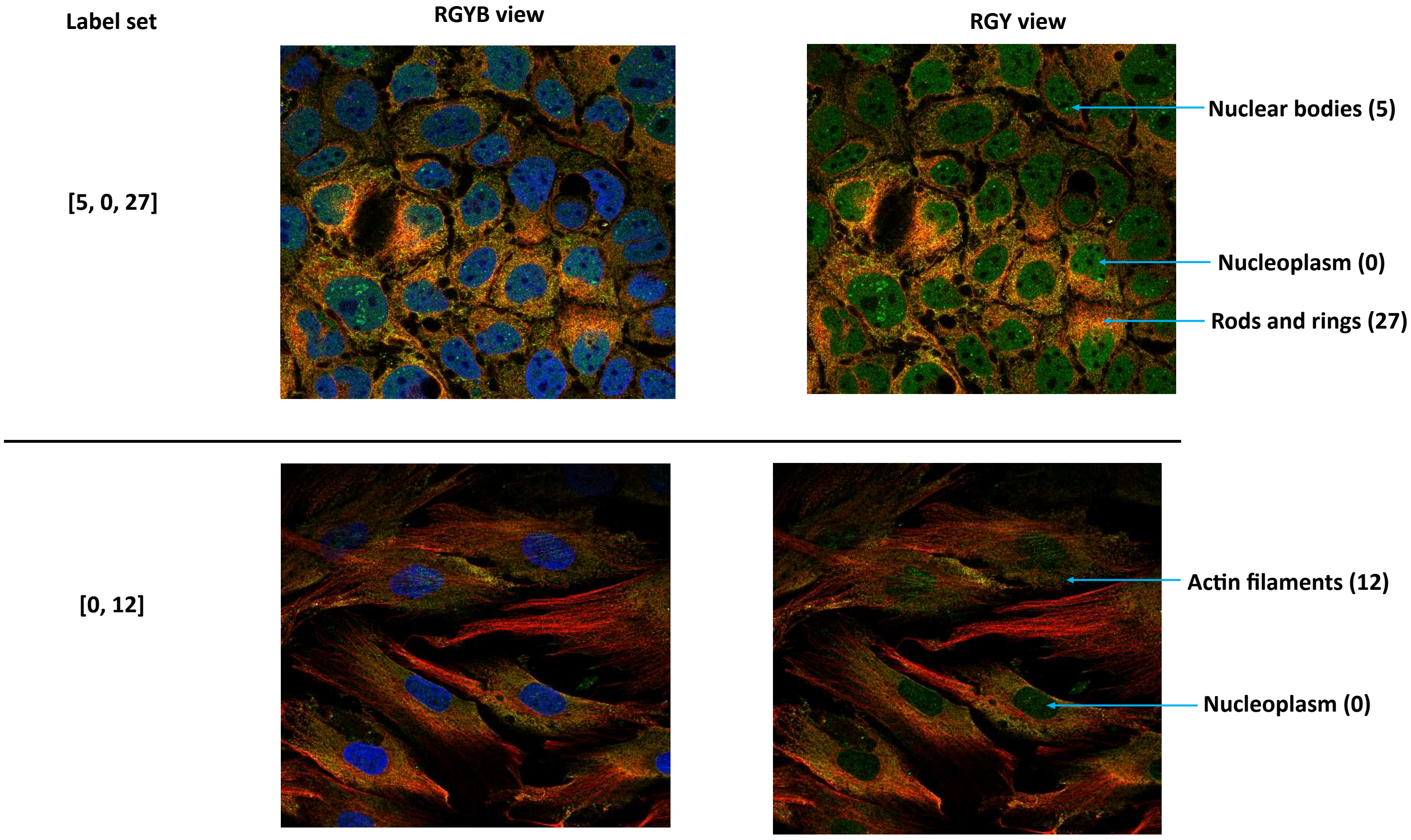


**Fig. 2.** Sample images showing multilocalised protein patterns in different cell lines that makes the protein subcellular localisation task challenging.

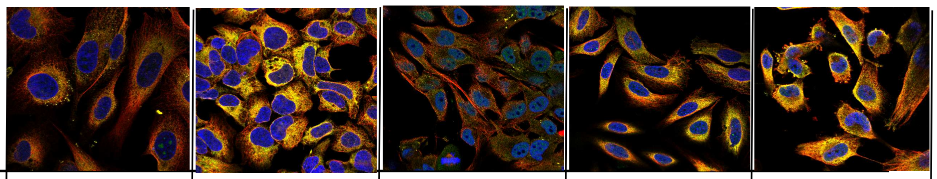

| | | | | | | |
|---|---|---|---|---|---|---|
| Original label set | | [0,8] | [2,7,8] | [0,15] | [0,5,15] | [23,27] |
| Predicted label set | Our method | [0,8] | [2,7,8] | [0,15] | [0,5,15] | [23,27] |
| | Wang, 2020 | [0,7] | [2,7] | [0] | [5] | [23] |

**Fig. 3.** Minority class sample images from TS1 demonstrate better performance of our method than Focal + Lovász loss (Wang, 2020).

#### 2 Details of applied metric learning

In metric learning, the definition of the positive and negative pairing is crucial, to make the model learn to distinguish between images. However, it is not straightforward in the multilabel environment. We explored triplet margin loss function (Ding *et al.*, 2015) with weighted sampling to evaluate metric learning, as unlike contrastive loss, triplet margin loss is able to tolerate some intraclass variance. Similar to contrastive loss function, the triplet margin loss function also aims to project similar images nearby and dissimilar images far away from each other. In order to achieve that, triplet margin loss builds triplets, which constitutes an anchor image, a positive image which is of the same class as the anchor, and a negative image which is of a different class from the anchor. Since protein subcellular localisation is a multilabel classification task of protein images with more than $500$ different label sets and high intraclass variance due to images from different phenotypes, we applied overlapping (an estimate of the number of common labels) of label sets as a criterion to mine the positive/negative pairs. In a minibatch, the image which has the maximum overlap of labels with the anchor image is considered positive, while the image pairs with the number of common labels as zero (specifically for minority classes) or one are considered as negative pairings.

Hyperparameter margin for triplet margin loss was set to $0.5$.
